# cancercelllines.org - a Novel Resource for Genomic Variants in Cancer Cell Lines

**DOI:** 10.1101/2023.12.12.571281

**Authors:** Rahel Paloots, Michael Baudis

## Abstract

Cancer cell lines are an important component in biological and medical research, enabling studies of cellular mechanisms as well as the development and testing of pharmaceuticals. Genomic alterations in cancer cell lines are widely studied as models for oncogenetic events and are represented in a wide range of primary resources. We have created a comprehensive, curated knowledge resource - cancercelllines.org - with the aim to enable easy access to genomic profiling data in cancer cell lines, curated from a variety of resources and integrating both copy number and single nucleotide variants (SNVs) data. We have gathered over 5,600 copy number profiles as well as SNV annotations for 16,000 cell lines and provide this data with mappings to the GRCh38 reference genome. Both genomic variations and associated curated metadata can be queried through the GA4GH Beacon v2 API and a graphical user interface with extensive data retrieval enabled using GA4GH data schemas under a permissive licensing scheme.

## 1 Introduction

Cancer genomes harbor numerous mutations that can serve in disease classification, clinical prediction as well as to identify targets for therapeutic interventions. These somatic mutations in cancers often occur in genes responsible for DNA replication or repair, enabling uncontrolled proliferation of cancer cells as well as introducing additional mutations to cancer genomes. Single nucleotide polymorphisms (SNPs) in these genes lead to altered functionality of the gene that is favorable for cancer progression. Another kind of variation often introduced to cancer genomes are structural variants where a large region in the chromosome is altered. Changes to the chromosome where large sequences are deleted or amplified are also referred to as copy number variants (CNVs). Oncogenes are genes endowing cell division functionalities also contribute cancer cell progression and additional copies of said gene further enhance cancer growth. Tumor suppressor genes are genes that induce cell death and ensure control mechanisms for correct replication processes, halting cancer progression. In addition to mutations in tumor suppressor genes that alter functionality, these genes are often deleted from cancer genomes. These copy number aberrations lead to deviations from normal diploid state of the genome and are often characteristic for cancer types. For instance, colorectal carcinomas are known to have a duplicated chromosome 13 that is characteristic for the disease [1, 2, 3].

While the molecular analysis of individual tumors provide enormous insights into molecular alterations affecting individual cases or disease types in general, testing of proposed mechanisms and design of pharmacological interventions require the observation of cells of individual tumor types under experimental conditions. These cell lines, representative for specific cancer types and carrying the respective molecular alterations, provide an ubiquitous tool set for foundational and translational cancer research settings. Cancer cell lines are usually obtained by extracting cells from individual tumors and are then cultivated under *in vitro* conditions. For certain applications - e.g. the generation of additional material for molecular testing - tumor derived cells are only expanded for a few divisions. However, the main application of *in vitro* systems lies in the establishment of immortalized cancer cell lines with the potential for long term storage and expansion in various experimental settings. Several public and commercial repositories provide thousands of such cell lines for many different cancer types, and some core sets of cell lines (e.g. NCI-60 [4]) are used in most screening systems for the definition of anti-neoplastic pharmaceuticals as well as in studies not directly related to cancer research.

Several databases have been created to collect and characterize cancer cell line data, such as Cellosaurus - a knowledge resource on cell lines [5]. While Cellosaurus includes extensive information on cell lines, such as their origin, age at collection, sex *etc*, other sources focus on the genomic data of cancer cell lines. Cancer Cell Line Encyclopedia (CCLE) is a database that characterises cancer cell line mutational data [6]. Genomic information relevant to cell lines is also included in curated resources such as ClinVar [7], *i.e*. information about effects of individual cancer-related mutations which also apply to cell lines. Another example of such resource is Progenetix, a reference resource for sample-specific genomic copy number variation (CNV) data of human cancers. In addition to primary cancers it also includes over 5,600 samples of human cancer cell line copy number variation profiles [8].

Here, we present our novel resource that focuses on cancer cell line variants. We have incorporated and curated data from various resources, including relevant metadata, allowing for numerous query types, *e.g*. searching by cancer type or variant position. Our resource also provides the unique opportunity to visualize available copy number and single nucleotide variants simultaneously.

## 2 MATERIALS AND METHODS

### 2.1 Variant resources

To create this database we collected data from the following resources: ClinVar [7], CCLE [6], NCIT [9], Cellosaurus [5] and Progenetix [8]. Specifically, human health related variants and associated metadata was derived from ClinVar while known variants from human cancer cell lines were obtained from CCLE. Existing information associated with the cell line and its source originate from Cellosaurus. NCIT cancer classification terms were used to standardize available diagnostic codes and represent diseases hierarchically. Progenetix was the source of our CN variants collection and also provided additional publications and geographical information on the cell lines. Moreover, Progenetix provides numerous visualization options particularly for copy number variations.

### 2.2 Variant import and mapping

CNV data for 5,754 copy number profiling experiments representing 2,163 individual call lines were imported from the Progenetix resource. Original data there had been generated from various platforms, predominantly genomic array experiments and including a core set of biomedical and technical metadata as described previously [8]. SNVs were mapped to Cellosaurus IDs and imported from ClinVar and CCLE resources. As for the CNV data we used the GRCh38 reference genome and applied a 0-based interbase coordinate system, in accordance with GA4GH recommendations and the GA4GH Variant Representation Specification (VRS [10]). For mutation data derived from the CCLE resource this process required the translation of the GRCh37 mapped data using a Python-based liftover tool [11].

#### 2.2.1 ClinVar

For the addition of cell line related genomic variants from ClinVar annotations we retrieved all sequence variation annotations from Cellosaurus. We then translated variants to appropriate format. For instance, Cellosaurus variant *STK11 p.Gln37Ter (c.109C>T)* was converted to ClinVar format (STK11):c.109C>T (p.Gln37Ter). Identifiers for the variants were then retrieved from (ClinVarVariationRelease_00-latest.xml, last accessed 2023-06-25). Available metadata for each variant were obtained by accessing ClinVar API, for example *https://eutils.ncbi.nlm.nih.gov/entrez/eutils/esummary.fcgi?db=clinvar&id=376334* for the aforementioned variant. Only variant locations for hg38 version were used. Additionally, MyVariant python package (version: 1.0.0) was used to add additional HGVS IDs to the variants.

#### 2.2.2 CCLE Mutations

Cancer cell lines in Cellosaurus dataset include a DepMap ID that is a unique identifier used for cell lines in CCLE data. We used these identifiers to retrieve CCLE mutational data. The mutation dataset was retrieved from CCLE “Downloads” site [12].

### 2.3 Cell line metadata

The essential part of the information associated with a cell line originates from Cellosaurus cell line resource [5]. Such information includes genome ancestry information, NCIT diagnostic classification of the cell line, age at collection *etc*. For the NCIT classification as well as for cell lines we apply a hierarchical entity representation. For the disease classification this corresponds to projection of all identified cell line disease codes to the NCIT “neoplasm” tree. For cell line identifiers we created a physical inheritance hierarchy tree in which derived cell lines are represented as children of the donor cell line and so forth and thereby allow to connect cell line originating from the same donor. Progenetix provides core metadata for CNV samples such as publications and provenance of the sample, enabling the geographical mapping of the samples [13].

### 2.4 Variant representation

All variants and associated metadata are represented using Beacon v2 default data model [14]. To accommodate to the requirements of cell line data, some elements of the schemas have been extended. We have used the following ontologies for mapping and standardizing our data: Human Ancestry Ontology (genome ancestry), Sequence ontology (variant effects), Phenotype And Trait Ontology (genotypic sex), NCIT (histological diagnoses), UBERON (anatomic topography), as well as ICD-O 3 for diagnoses (M) and topographic site (T).

### 2.5 Front-end and API

The *cancercelllines.org* resource is built from software stack incorporating a MongoDB database back-end, a middleware and API stack from the *bycon* and *byconaut* Python packages and the *cancercelllines-web* React-based Javascript front-end, served through an Apache webserver set-up. Here, the *bycon* software implements a Beacon v2 [14] API and the front-end interacts with this through Beacon conform requests and parsing of the standard Beacon JSON responses. For instance, a search for breast cancer cell lines with genomic variants in the TP53 gene locus would consist of a genomic range query (chromosome, base positions of TP53 start and end coordinates, with optional type or variant base such as “SO:0001059” for a sequence alteration) together with a disease code for breast carcinomas (*e.g*. “NCIT:C4872 - Breast Carcinoma”) which is provided as a “filter” to the Beacon API. In our implementation, primary Beacon v2 response will consist of a “count” response for the “biosample” entry type (*i.e*. provide the number of matched samples) with embedded “handover” links for the retrieval of the individual samples and genomic variations through asynchronous https requests, again using the Beacon v2 protocol.

## 3 RESULTS

### 3.1 Collection of human cancer cell lines

After mapping of available variants from Progenetix (copy number variants), CCLE and ClinVar (SNVs), we created entries for 16,000 unique cell lines from 400 different cancer types according to the NCIT classification. More than 15,000 of these cell lines have mapped variant(s) from ClinVar, over a 1,000 cell lines have one or more known SNVs for the cell line from CCLE and more than 2,000 cancer cell lines have available CNV profiles processed as part of the Progenetix data pipeline[8]. Table 1 shows some of the most common features of donors. In total, we have data from 4,214 donors, of those the genotypic sex is known for 1,882 male and 1,486 female samples. The average age of the donor is 50, with the minimum age being under 1 year old and 94 years old is the oldest. Ages of fetal samples were not included. The most frequent cancer type for all mapped cell line samples is melanoma (NCIT:C3224), followed by glioblastoma and lung small cell carcinoma. Interestingly, in all cases except female reproductive system carcinomas, number of male donors per diagnosis is higher than female donors.

**Table 1:**
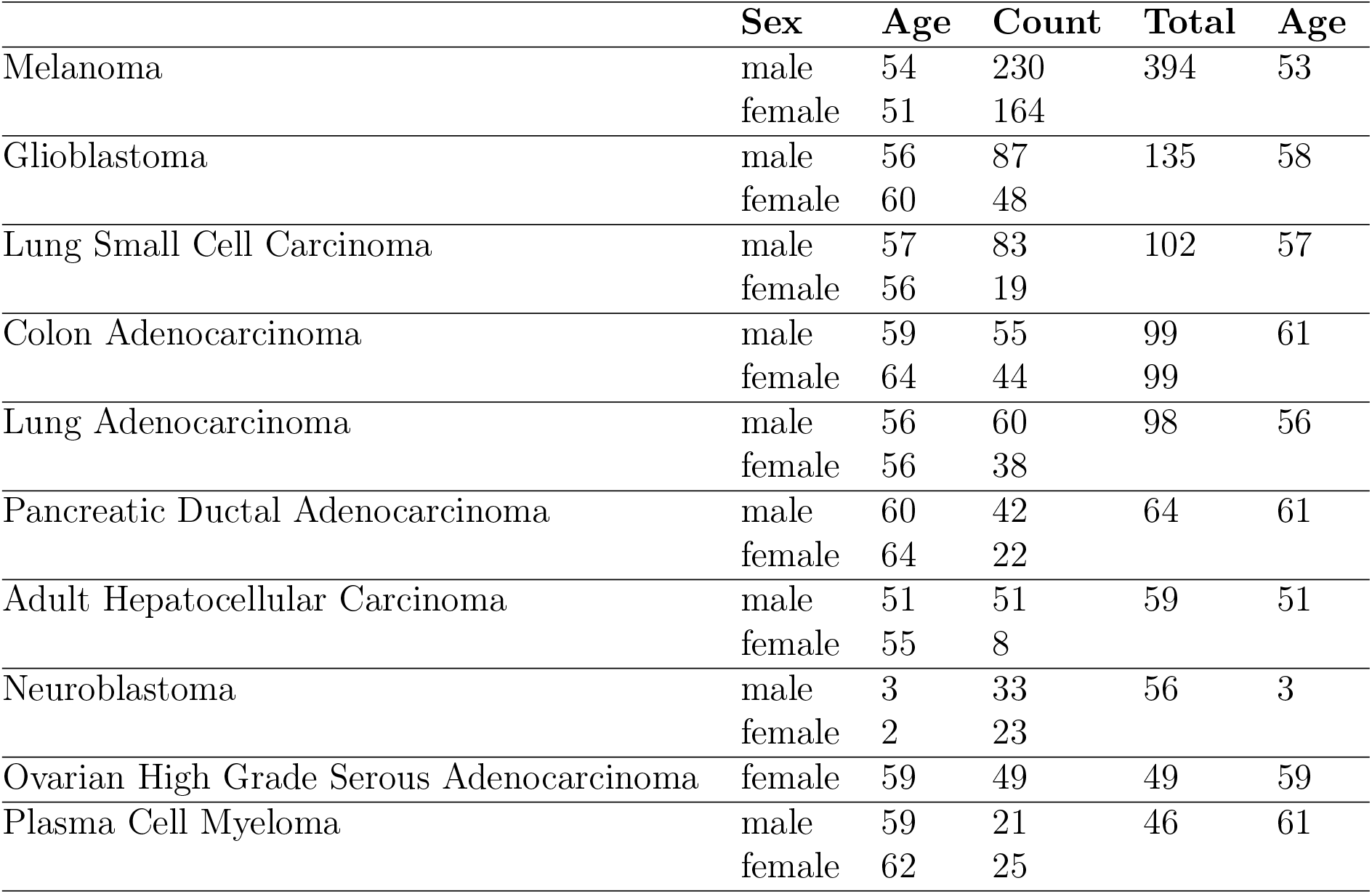
Cancer Cell Line Donor Statistics.

Hierarchical information on cancer cell lines can be found under “Cell Line Listings” on the left. There, the root level of each cell line is shown, child levels can be accessed by expanding. The search box in “Cell Line Listings” also allows for hierarchical queries of cell lines. The resulting landing page displays known metadata about the donor of the cell line as well as known parent and child terms.

Figure 1A illustrates the results for the first human cell line continuous established-HeLa [15]. HeLa, a cervical carcinoma cell line, was created in 1951 and the name was derived from the patient’s initials [16, 15]. Even today, 70 years later, HeLa is still one of the most widely used cell lines.

**Figure 1:**
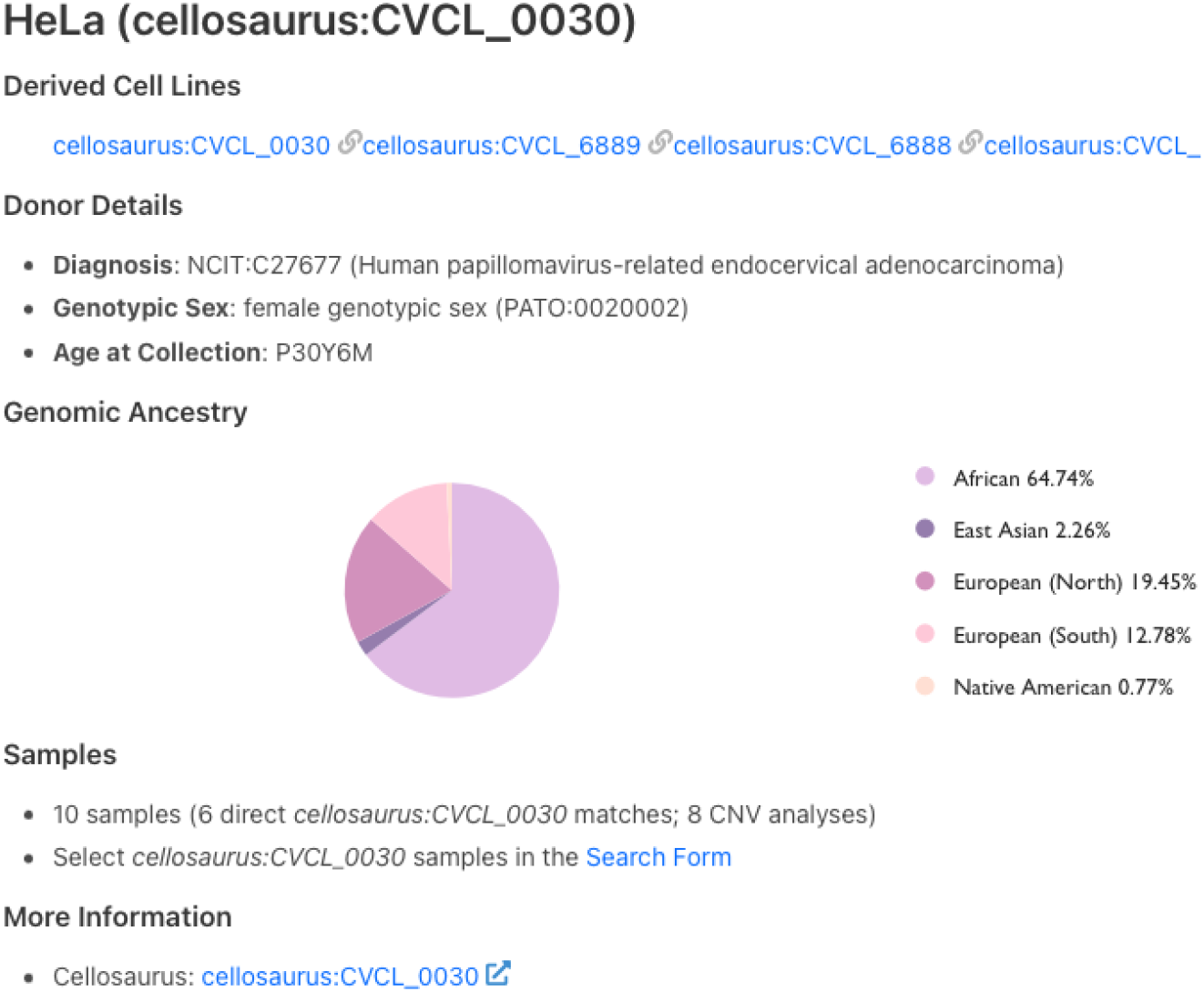
Cell line details page for *HeLa*. Listed on this page are derived cell lines and information on the cell line donor. The count of associated samples and link to “Search Form” are also shown. The last link on the page redirects to cell line page on Cellosaurus.

All mapped cell lines have a Cellosaurus ID and include metadata such as NCIT disease code associated with said cell line as well as genotypic sex of the material and age at collection. Additionally, for some cell lines genome ancestry data is also available and represented according to the Hancestro (Human Ancestry Ontology) model (Fig. 1). CNV frequency plots for the samples of cell line of interest as well as available child terms are shown, followed by mapped SNVs in the annotated variants section.

Moreover, information extraction results for annotated cancer cell line gene information is located under “Literature Derived Contextual Information”. More information on this can be found at Smith *et al*, 2023 [17].

### 3.2 Cancer cell line CNV profiles

The CNV data of cancer cell lines originate from Progenetix database where samples related to a cell line have previously been identified from open-source repositories such as Gene Expression Omnibus (GEO) or from data provided with original publications [8]. Figure 2 shows ratios of samples per NCIT diagnostic code of cancer cell lines and their origins in Progenetix. Most frequent cancer type among CNV samples in both cell lines and tumors is ductal breast carcinoma. The large number of breast carcinoma samples could be explained by the large breast cancer detection campaigns that have been implemented worldwide. Unexpectedly, melanomas are underrepresented among primary tumors compared to the sample number in cell lines. A disproportionate number of melanoma samples originates from some studies with a high number of melanoma cell line samples. For example, over 100 cutaneous melanoma samples were retrieved from a comparative study of copy number profiles [18]. A well portrayed origin group is chronic lymphocytic leukemia that is represented by only 3 cell line samples. One possible explanation could be that slowly progressing cancer types do not acclimate well to *in vitro* environment.

**Figure 2:**
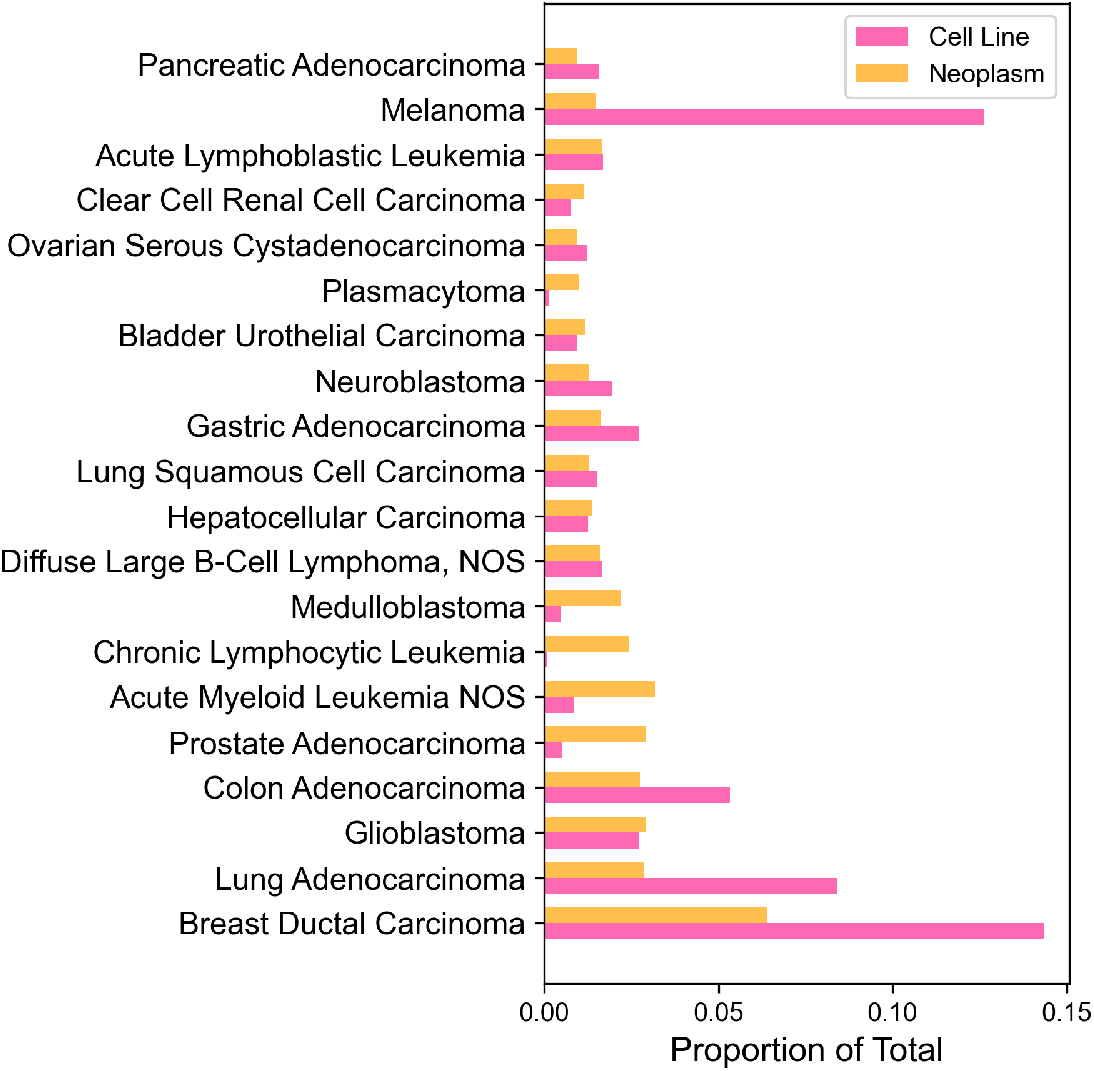
Comparison of sample numbers in cell lines and their origins for the most common cancer types. 20 most common cancer types (by the number of sample count, excluding “Unspecified Tissue” samples) were picked from Progenetix. Cancer types without any cell lines were excluded as well. Horizontal bars represent the proportion of total sample count for each cancer type. Cell line ratios are shown in pink and origin ratios are shown in orange.

It has been shown that cancer cell lines indeed exhibit similar CNV profiles to their origins but have a higher number of mutations [19]. Unfortunately, many of the widely used cancer cell lines have been found to be either contaminated or misidentified [20]. For instance, cell line MDA-MB-435 was thought to be a breast cancer cell line but was instead found to be originating from melanoma [21]. Figure 3 demonstrates cell line MDA-MB-435 compared to ductal breast carcinoma and amelanotic melanoma CNV profiles. By combining the data available in Progenetix and cancercelllines.org, we show that indeed MDA-MB-435 is more similar to amelanotic melanoma.

**Figure 3:**
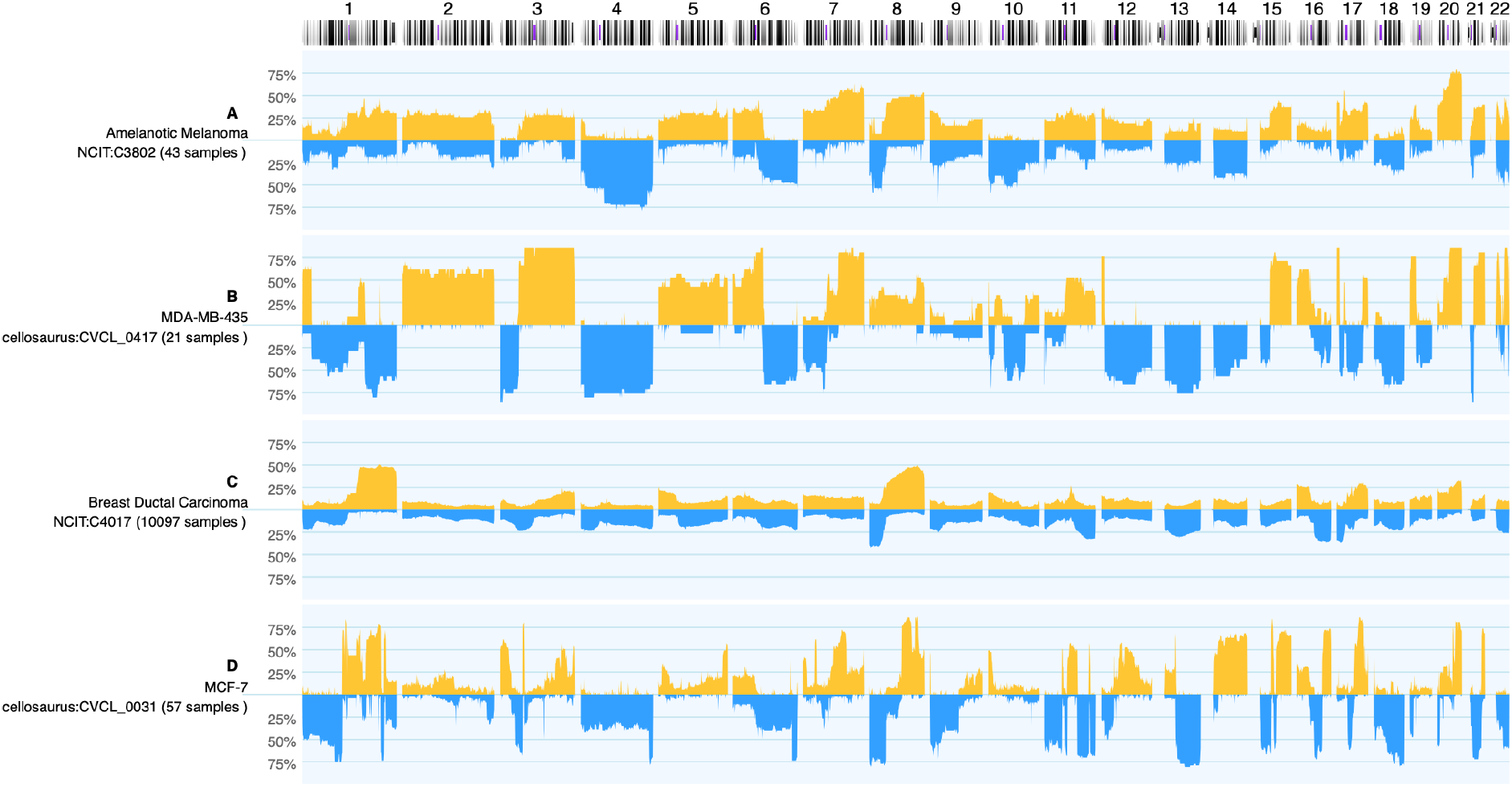
Genomic copy number variation frequencies comparing cancer-type specific profiles to those from selected cell lines for copy number gains (up, yellow) and losses (down, blue; 100% - CNV observed in all samples). While A and C display the summary data from 43 Amelanotic Melanomas (NCIT:C3802) and 10,254 Ductal Breast Carcinomas, respectively, panel C and D show summary profiles of cell lines MDA-MB-435 (from 21 instances) and MCF-7 (57 analyses). Although both cell lines were originally classified as *Breast Carcinoma*, the CNV pattern of MDA-MB-435 shows an intriguing similarity to aberrations common in Amelanotic Melanomas (e.g. +2, +3q, +5, +6p/-6q…). As of note, while the CNV frequency plots are influenced by the expected genomic heterogeneity of tumor samples the *in principle* expected genomic homogeneity of cell lines (*i.e*. either 100 or 0%) can be perturbed by genomic instability and leading to inter-sample variations as well as experimental conditions.

### 3.3 SNVs of cancer cell lines

To curate the SNVs in cancer cell lines and their effect on health, we mapped known cell line SNVs to ClinVar variants and pulled cancer cell lines from the CCLE mutations dataset. Table 2 shows the number of resulting variants from ClinVar and CCLE resources. Since ClinVar is a resource for variants related to human health, the number of distinct variants is lower than in CCLE that includes all variants from a set of cell lines. While CCLE only includes around 1,000 well characterized cancer cell lines, known human health-related variants have been found in over 15,000 cancer cell line entities. Surprisingly, the most commonly mutated gene in CCLE dataset is TTN, a gene responsible for producing titin protein that is the largest human protein and is a structural sarcomeric protein. Expectedly, the most common gene in ClinVar dataset is TP53, a tumor suppressor gene that is one of the most frequently mutated genes in cancer [22].

**Table 2:** SNV Statistics.

|  | ClinVar | CCLE mutations |
| --- | --- | --- |
| Unique variants | 1,246 | 23,947 |
| Total number of variants | 30,960 | 1,013,244 |
| Number of cell lines | 15,750 | 1,292 |
| Most frequent gene | TP53 | TTN |
| Number of genes | 144 | 18,739 |

### 3.4 Use cases

#### 3.4.1 CNV profiles

Our resource enables finding CNV profiles for cancer cell lines of interest, including the option to automatically include instances of derived cell lines. The query can be performed under “Search Cell Lines” section and subsequently using cell line name or Cellosaurus ID to search. For example, HeLa could be queried by typing “HeLa” or “CVCL_0030” into the ID field. Search query can also be executed by the diagnostic code of the cell line (NCIT) in the “Cancer Classification(s)” field. By default, all child terms found for the cell line (or diagnostic code) will be included. This can be changed under “Include Child Terms” field. The resulting landing page will show resulting CNV frequency plot with options to list existing biosamples, to see where these samples were from geographically and also to list the known annotated variants for these cell lines. Additional visualization options can be found under “Visualization options”. In addition to CNV frequency plots, this enables clustered view of the samples.

#### 3.4.2 SNV data

To access our SNV data, cell line on interest can be queried in “Search Cell Lines” like for CNV samples but additional field needs to be entered under Query by Position, Variant Type: SO:0001059 (any sequence alteration - SNV, INDEL). The resulting matched SNVs can be found under “Variants” section (Fig 4A). There, variants are listed and can be sorted by the field of interest. “Digest” shows the genomic location and the affected nucleotides of the variant and “Gene” represents the affected gene. Pathogenicity refers to known clinical impact of the variant from ClinVar annotations. Variant effect shows the effect of the mutation, according to sequence ontology.

**Figure 4:**
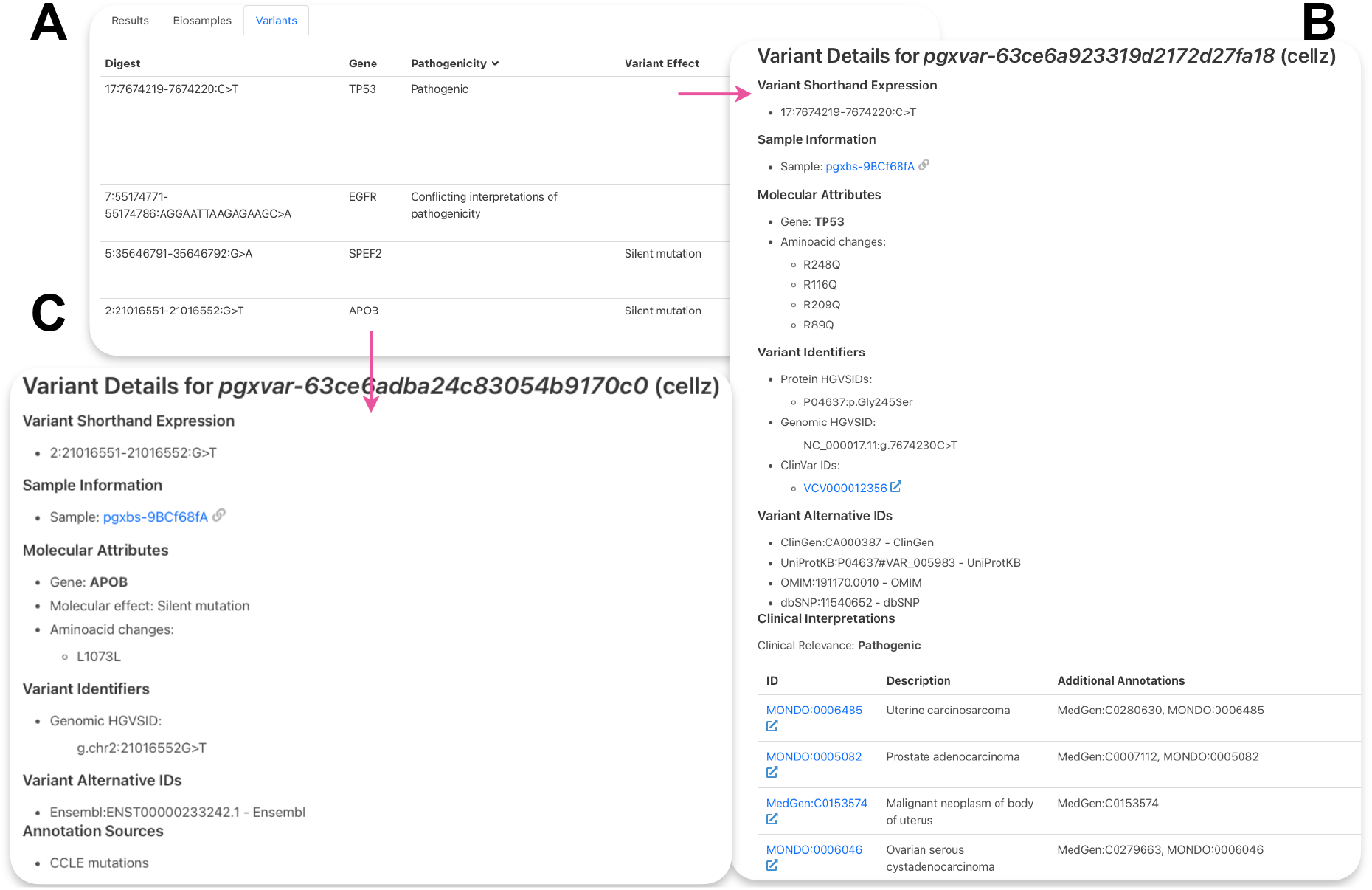
Lung adenocarcinoma cell line PC-9 SNVs. A - Table of resulting variants for PC-9. “Digest” shows the genomic location and the affected nucleotides of the variant, “Gene” - the affected gene, “Pathogenicity” - reported effect of the variant on human health, “Variant Effect” - effect of the variant on the gene product. B - example of a ClinVar variant. C - example of a CCLE mutation.

Clicking on the variant ID leads to variant page (Fig 4B and 4C). Figure 4B shows the results for a variant in TP53 from ClinVar. Shown are available variant HGVS identifiers, ClinVar identifiers as well as alternative IDs for this variant from other resources. Under “Clinical Interpretations”, disease ontologies are listed by ID and descriptions are provided on the left. Clicking on the ID of interest will redirect to the disease ontology page. Figure 4C shows available mutation data for a variant in the APOB gene. ClinVar is a database for phenotypic health-related variants, therefore each variant includes more information compared to CCLE, that only shows cancer cell line specific variant information. Information about the molecular attributes of this mutations can be found for CCLE variants such as amincoacid changes and molecular effect. Some genomic HGVS identifiers are also available for CCLE variants. More information is included in a detailed user guide (Supplementary Materials).

## 4 DISCUSSION

Cancer cell lines are important model systems in many areas of biomedical research. While the knowledge about their genomic variations represents an essential component for their effective and accurate utilization, this information is dispersed in different types of databases and repositories. Here, we have presented *cancercelllines.org*, a website and knowledge resource with a comprehensive collection of curated cancer cell line genome variants. In this database we have included a large collection of annotated sequence variants and generated copy number profiling data as well as curated metadata including identifier-based links to external donor repositories and information resources. Importantly, cell line entities are linked hierarchically according to their provenance thereby facilitating analyses of mutational dynamics as well help with the identification of labeling inconsistencies. While various excellent resources such as COSMIC [23] and CCLE [6] contain data about genomic variants in cancer cell lines, our resource offers a unique, comprehensive functionality to assess genomic data combined from various resources, including a large, unique set of genome-wide CNV profiling data.

Cancer cell lines can be used in different fields in life sciences but predominantly serve to study disease mechanisms of cancers and evaluate potential targets of therapeutic interference. The use of these model systems in conjunction with genome profiling data from native tumor samples can be advantageous to select cell lines for *in vitro* experiments matching the tumor types of interest, potentially beyond the confinements of diagnostic classifications. The comparison of CNV profiling data between cancer cell lines and native tumor samples may provide a new avenue for the use of cancer cell line models in “matched genomics” scenarios. The integration of cancer cell line data with data from the Progenetix resource - facilitated through common frameworks, annotation standards, query and visualization methods - enables both the visual identification of similarities in data patterns (*cf*. figure 3), as well as the retrieval of standardized data for offline analyses.

Data discovery and retrieval in *cancercelllines.org* is enabled through the use of the “Beacon” API, a standard of the Global Alliance for Genomics and Health (GA4GH), and associated schemas for genomic as well as biomedical and technical metadata. Importantly, the support of these standards in an open access data setting allows the integration of *cancercelllines.org* into federated data discovery scenarios [24] where each resource provides complementary data under a common access protocol.

## Supporting information

User Guide

## 5 ACKNOWLEDGEMENTS

We would like to thank Prof. Dr. Amos Bairoch, the founder and director of the Cellosaurus resource for his extensive support and advice. The work was supported through ELIXIR as part of the “Beacon and beyond - Implementation-driven standards and protocols for CNV discovery and data exchange” project.

## 5.0.1 Conflict of interest statement

None declared.

## 6 Supplementary Information

We have provided the following supplementary files:

- User guide (PDF)

## Notes

https://cancercelllines.org

### Competing Interest Statement

The authors have declared no competing interest.

