## Supplementary material for "cancercelllines.org - a Novel Resource for Genomic Variants in Cancer Cell Lines": User Guide

### cancercelllines.org user manual


General 1

Cell Lines Listing 2

Search Cell Lines 4

CNV Profiles by Cancer Type 12


#### General

On the front page you can find some background information on our database and a randomly created cancer cell line CNV frequency plot where amplifications are shown in yellow and deletions in blue. The left panel shows different query options.

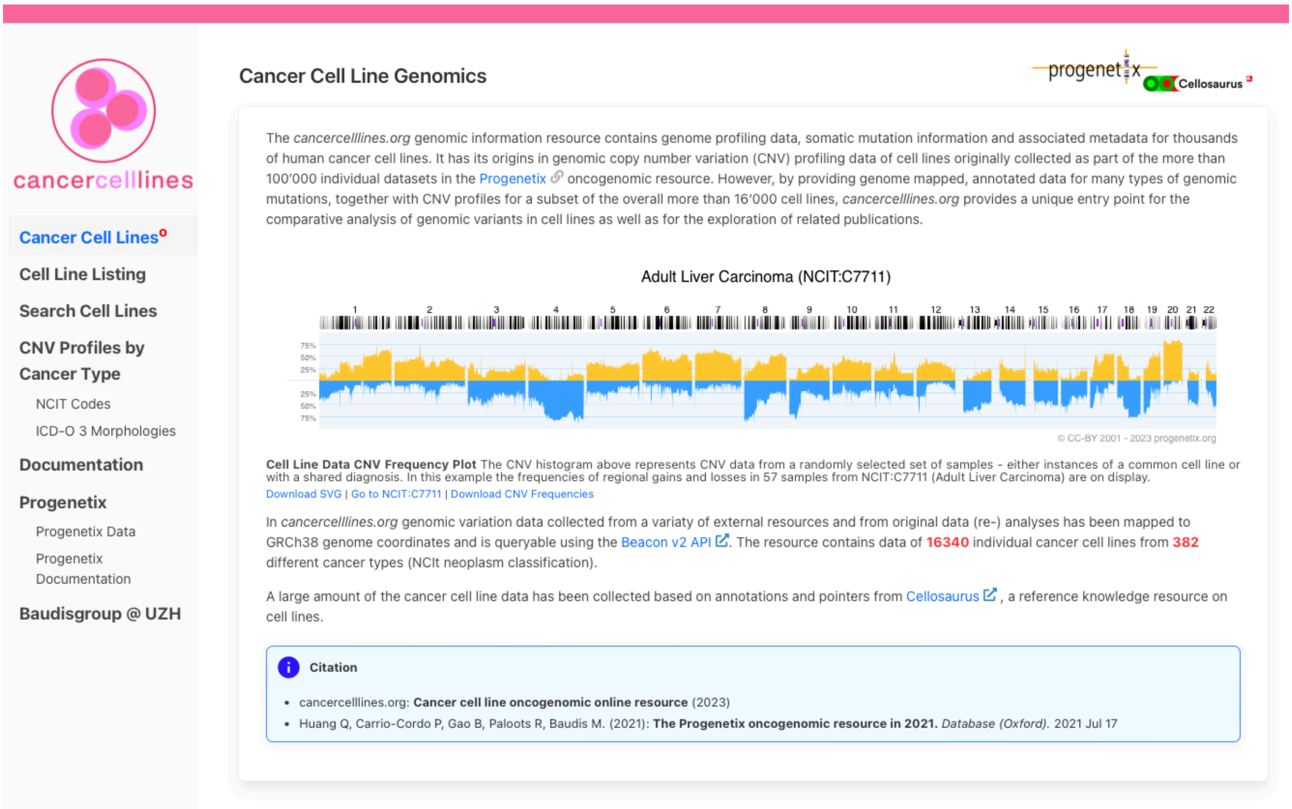


**Cell Line Listings** - find all available cancer cell lines represented hierarchically. Search box allows for the query of cell line of interest. Returns info on cell line origins, hierarchies, variants and cell line gene information extraction.
**Search Cell Lines** - general search box for variant queries, cell line specific queries and queries by cancer type.

**CNV profiles by Cancer Type** - get cancer cell line CNV profiles for cancer type of interest. Get hierarchical cancer types by NCIT or ICDO classifications.

### Cell Lines Listing

#### 1. Search Form

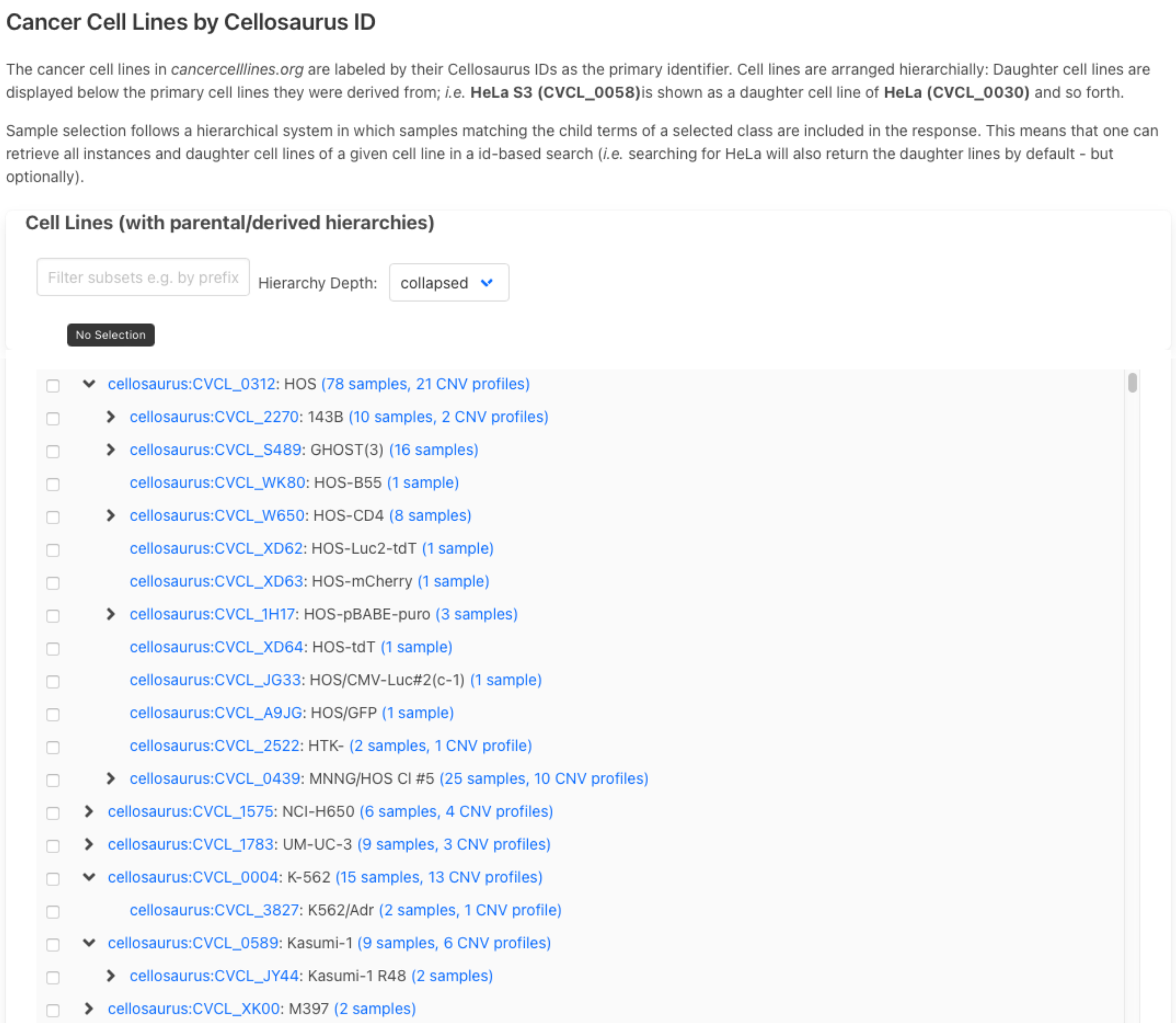


On "Cell Lines Listing" page, available cancer cell lines are listed and child terms can be expanded by clicking on the ">". You can also search for a cell line either by name or cellosaurus ID in the search box. Numbers indicate available biosamples and CNV profiles available for cell line of interest. By clicking on cell line you will be directed to cell line results page.

#### 2. Cell Line Page

### K-562 (cellosaurus:CVCL_0004)

##### Derived Cell Lines

cellosaurus:CVCL_3827 cellosaurus:CVCL_0004

##### Donor Details

- **Diagnosis**: NCIT:C9110 (Blast phase chronic myelogenous leukemia, BCR-ABL1 positive)
- **Genotypic Sex**: female genotypic sex (PATO:0020002)
- **Age at Collection**: P53Y

##### Genomic Ancestry

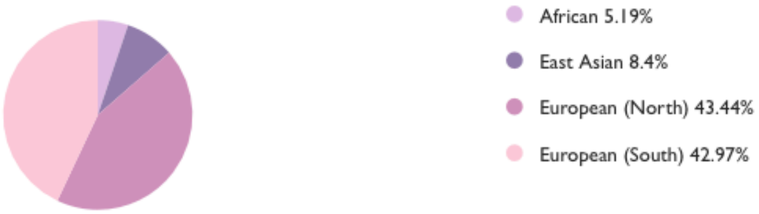


##### Samples

- 15 samples (13 direct *cellosaurus:CVCL_0004* matches; 13 CNV analyses)
- Select *cellosaurus:CVCL_0004* samples in the Search Form

##### More Information

- Cellosaurus: cellosaurus:CVCL_0004

Cell line results page first gives information on the cell line. First, derived and parental cell lines are listed. Then information on the donor individual is listed including diagnosis, sex and age. Genomic ancestry is also available for a selection of cell lines. Below is a section on sample numbers and further search options. A link to Cellosaurus cell line knowledge resource is also provided for more information.

**Annotated Variants for cellosaurus:CVCL_0004**

| Digest | Gene ^ | Pathogenicity | Variant Effect | Variant Instances |
|---|---|---|---|---|
| 17:68932401-68932403:->A | ABCA8 | | Frameshift variant | V: pgxvar-63ce6acda24c83054b8bbc02<br>B: pgxbs-c43B6e8E |
| 7:48278880-48278881:C>A | ABCA13 | | Missense variant | V: pgxvar-63ce6acda24c83054b8bbcd3<br>B: pgxbs-c43B6e8E |
| 7:87409347-87409348:G>A | ABCB4 | | Missense variant | V: pgxvar-63ce6acda24c83054b8bbcd7<br>B: pgxbs-c43B6e8E |
| 11:34356773-34356774:G>T | ABTB2 | | Silent mutation | V: pgxvar-63ce6acda24c83054b8bbb90<br>B: pgxbs-c43B6e8E |
| 2:210205750-210205751:C>A | ACADL | | Missense variant | V: pgxvar-63ce6acda24c83054b8bbc5e<br>B: pgxbs-c43B6e8E |
| 5:33534833-33534834:G>C | ADAMTS12 | | Splice region variant | V: pgxvar-63ce6acda24c83054b8bbca0<br>B: pgxbs-c43B6e8E |
| 11:130449684-130449685:C>A | ADAMTS15 | | Missense variant | V: pgxvar-63ce6acda24c83054b8bbba8<br>B: pgxbs-c43B6e8E |
| 5:5209159-5209161:->A | ADAMTS16 | | Frameshift variant | V: pgxvar-63ce6acda24c83054b8bbca1<br>B: pgxbs-c43B6e8E |
| 6:143433897-143433898:A>C | ADAT2 | | Missense variant | V: pgxvar-63ce6acda24c83054b8bbcda<br>B: pgxbs-c43B6e8E |
| 4:7773002-7773003:C>A | AFAP1 | | Missense variant | V: pgxvar-63ce6acda24c83054b8bbc90<br>B: pgxbs-c43B6e8E |

« ‹ › » Page **1 of 48**

The following section shows annotated single nucleotide variants for said cell line. Variants can be ordered by gene, for example, by clicking “Gene” on the table header. Variants shown in this section include derived variants from parental cell lines.

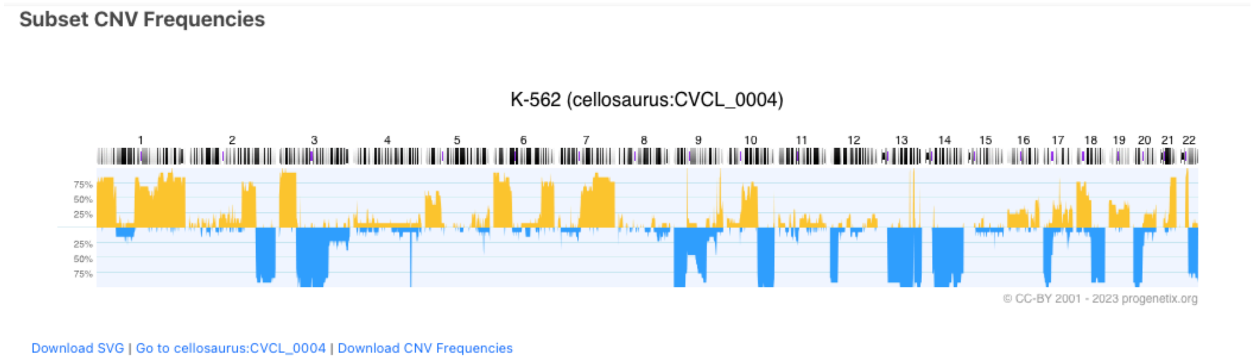


Subset CNV frequencies are visualized in the next section. Links below frequency plot direct to downloading the plot in SVG format as well as to the search page.

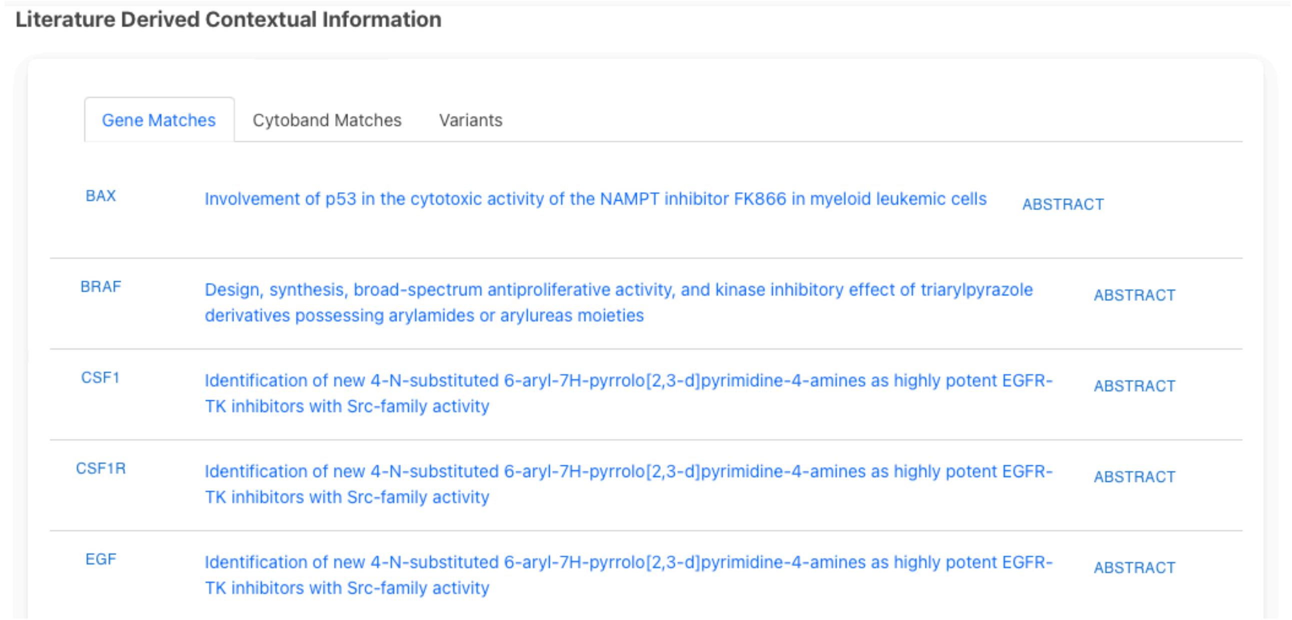
Literature Derived Contextual Information

Gene Matches | Cytoband Matches | Variants

| | | |
|---|---|---|
| BAX | Involvement of p53 in the cytotoxic activity of the NAMPT inhibitor FK866 in myeloid leukemic cells | ABSTRACT |
| BRAF | Design, synthesis, broad-spectrum antiproliferative activity, and kinase inhibitory effect of triarylpyrazole derivatives possessing arylamides or arylureas moieties | ABSTRACT |
| CSF1 | Identification of new 4-N-substituted 6-aryl-7H-pyrrolo[2,3-d]pyrimidine-4-amines as highly potent EGFR-TK inhibitors with Src-family activity | ABSTRACT |
| CSF1R | Identification of new 4-N-substituted 6-aryl-7H-pyrrolo[2,3-d]pyrimidine-4-amines as highly potent EGFR-TK inhibitors with Src-family activity | ABSTRACT |
| EGF | Identification of new 4-N-substituted 6-aryl-7H-pyrrolo[2,3-d]pyrimidine-4-amines as highly potent EGFR-TK inhibitors with Src-family activity | ABSTRACT |

The last section of the cell line results page shows literature derived gene information on the cell lines. Click on the gene name to visualize it on the frequency plot. Clicking on the article title redirects you to the article page on PubMed. Abstract of the paper will be expanded upon left-click.

### Search Cell Lines

You can also search for individual cell line by the name or ID or search for cell lines that match the diagnosis of interest (*eg* NCIT code) or search for all cell lines that harbour a specific variant.

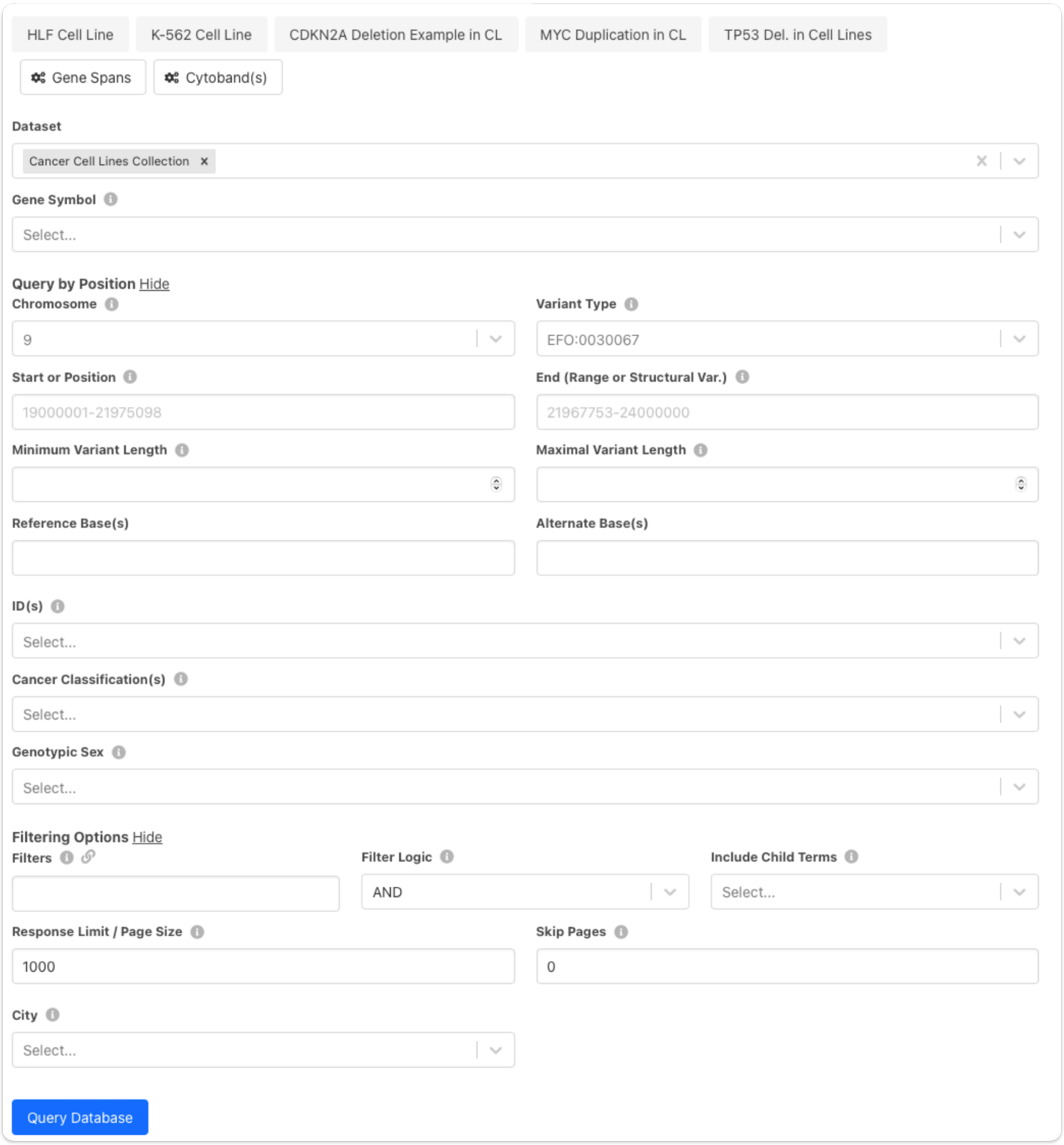


- The top row of the search fields offers some example queries, for both cell lines and positional queries.
- You can use *Gene Spans* and *Cytobands* buttons to limit your results to a range of a gene of interest or a specific cytoband.
- Default dataset is Cancer Cell Lines Collection but Progenetix collection is also available to enable queries with tumor samples
- *Gene Symbol* - search for a variant in a gene
- *Query by Position* - show fields to enable search by position
- *Chromosome* - enter chromosome of interest. Refseq IDs are also accepted.
- *Variant Type* - specify variant type by EFO term. This field can be used to receive available SNVs.

- *Start or Position* - start of sequence variation of 1-based genomic positions
- *End* - end of sequence variation of 1-based genomic positions
- *Minimum/Maximum variant length* - for queries with pre-defined variant lengths
- *Reference/Alternate Base(s)* - enter reference and altered nucleotide bases
- *IDs* - this field can be used to query cell lines by their IDs or to search for results of different PubMed IDs. Cellosaurus IDs or cell line names (Cellosaurus nomenclature) are accepted as input for cell lines.
- *Cancer Classification(s)* - use NCIT or ICDO disease codes to search for cell lines of the diagnostic code.
- *Genotypic Sex* - search for male or female cell line samples.
- *Filtering options* - additional filters to limit the number of search result, define filter logic and include child terms.
- *Filters* - option to add additional free comma separated filters *e.g.* PMID, CVCL etc.
- *Filter logic* - and/or
- *Include Child Terms* - includes available child terms or only shows exact matches. Can be applied for cell lines and diagnostic codes as well.
- *Response Limit/Page Size* - default is 1000, can be increased or decreased
- *Skip Pages* - select number of pages to be skipped
- *City* - search samples by the annotated city

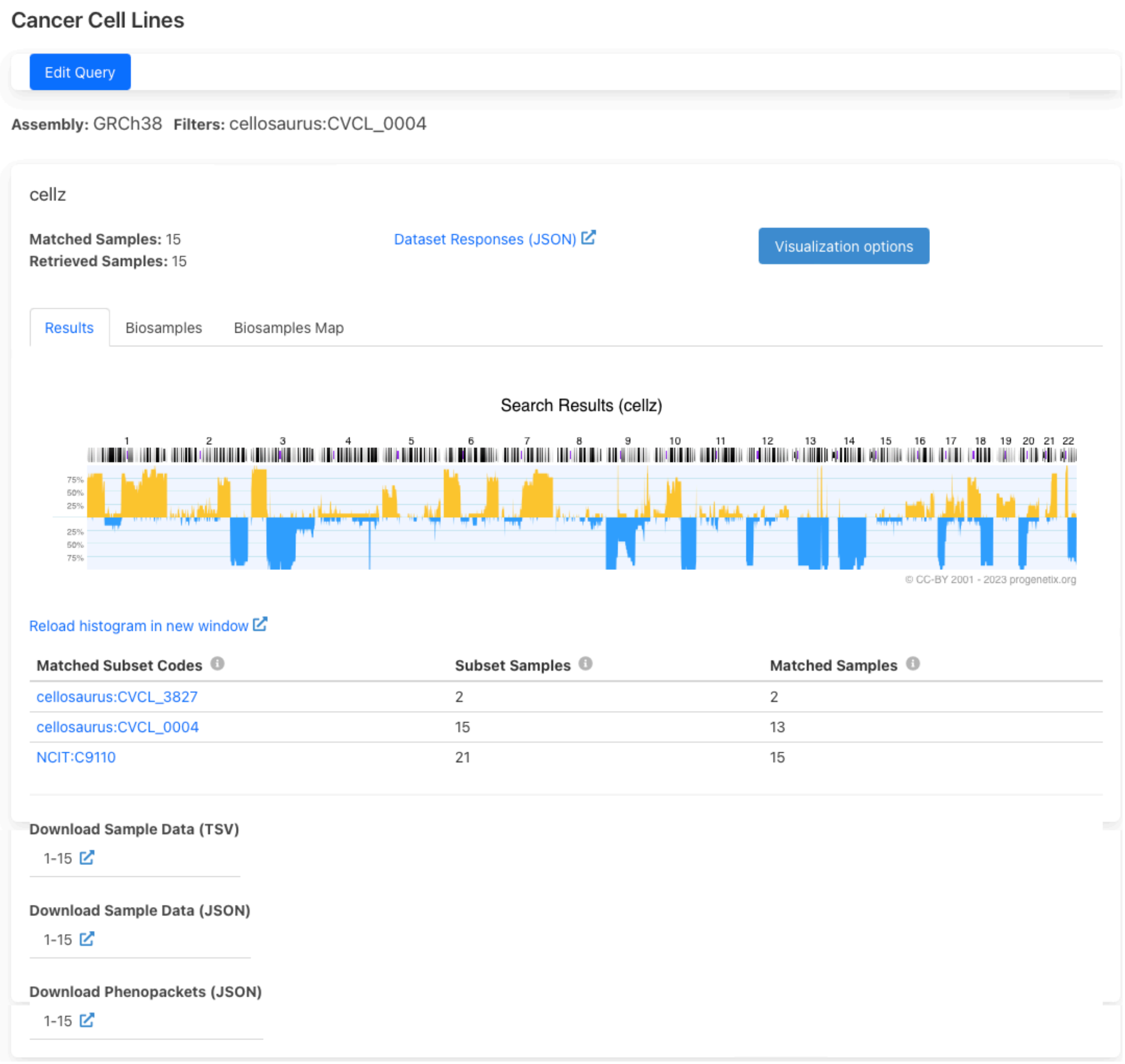

### Search Results: Cell Line Query

Here we show the search results for an example query listed in the search form - K-562 cell line. All data in our database is uses GRCh38 assembly version. Used filters, in this case Cellosaurus ID for K-562 are shown on top. Cellz - an internal short name for the cancer cell lines dataset. Dataset responses can be viewed in a JSON format. In the results tab, the frequency plot for query result is shown. On the last page, you can find a table of matched subsets and sample numbers. These data can also be downloaded in TSV or JSON format.

### Biosamples

##### Sample Details for *pgxbs-kftvipwj*

**Description**

Chronic myelogenous leukemia [cell line K-562]

**Diagnostic Classifications**

- Blast phase chronic myelogenous leukemia, BCR-ABL1 positive: NCIT:C9110
- Chronic myelogenous leukemia, BCR/ABL positive: pgx:icdom-98753
- Bone marrow: pgx:icdot-C42.1
- bone marrow: UBERON:0002371

**Cell Line Info**

- Instance of cellosaurus:CVCL_0004

**Donor Details**

- **Diagnosis**: NCIT:C9110 (Blast phase chronic myelogenous leukemia, BCR-ABL1 positive)
- **Genotypic Sex**: female genotypic sex (PATO:0020002)
- **Age at Collection**: P53Y

**Genomic Ancestry**

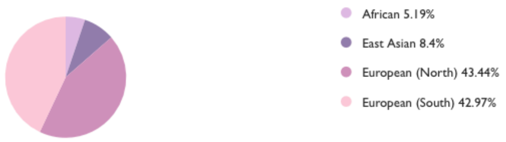


**Provenance**

- Origin: Boston, United States

**External References**

- Garraway LA, Widlund HR et al. (2005): Integrative genomic analyses identify MITF as...: PMID:16001072
- K-562: cellosaurus:CVCL_0004
- K-562: geo:GSM50195
- Affymetrix 100K SNP array data-: geo:GSE2520
- Affymetrix GeneChip Human Mapping 50K Hind - prerelease: geo:GPL2014

**CNV Plot**

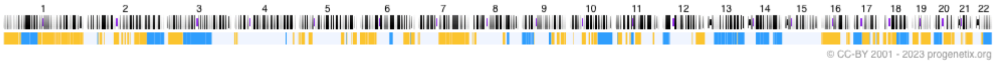


**Download**

- Sample data as Beacon JSON
- Sample data as Beacon Phenopacket JSON
- Sample variants as Beacon JSON
- Sample variants as Progenetix .pgxseg file
- Sample variants as (experimental) VCF 4.4 file

**Raw Data**

⇒ click to show/hide

Clicking on “Biosample ID” on the Biosamples tab will lead to the sample page. There info about the cell line origin is shown - same as in Cell Line Page. Additional reference for the sample are provided and CNV plot of the sample is shown (when available).

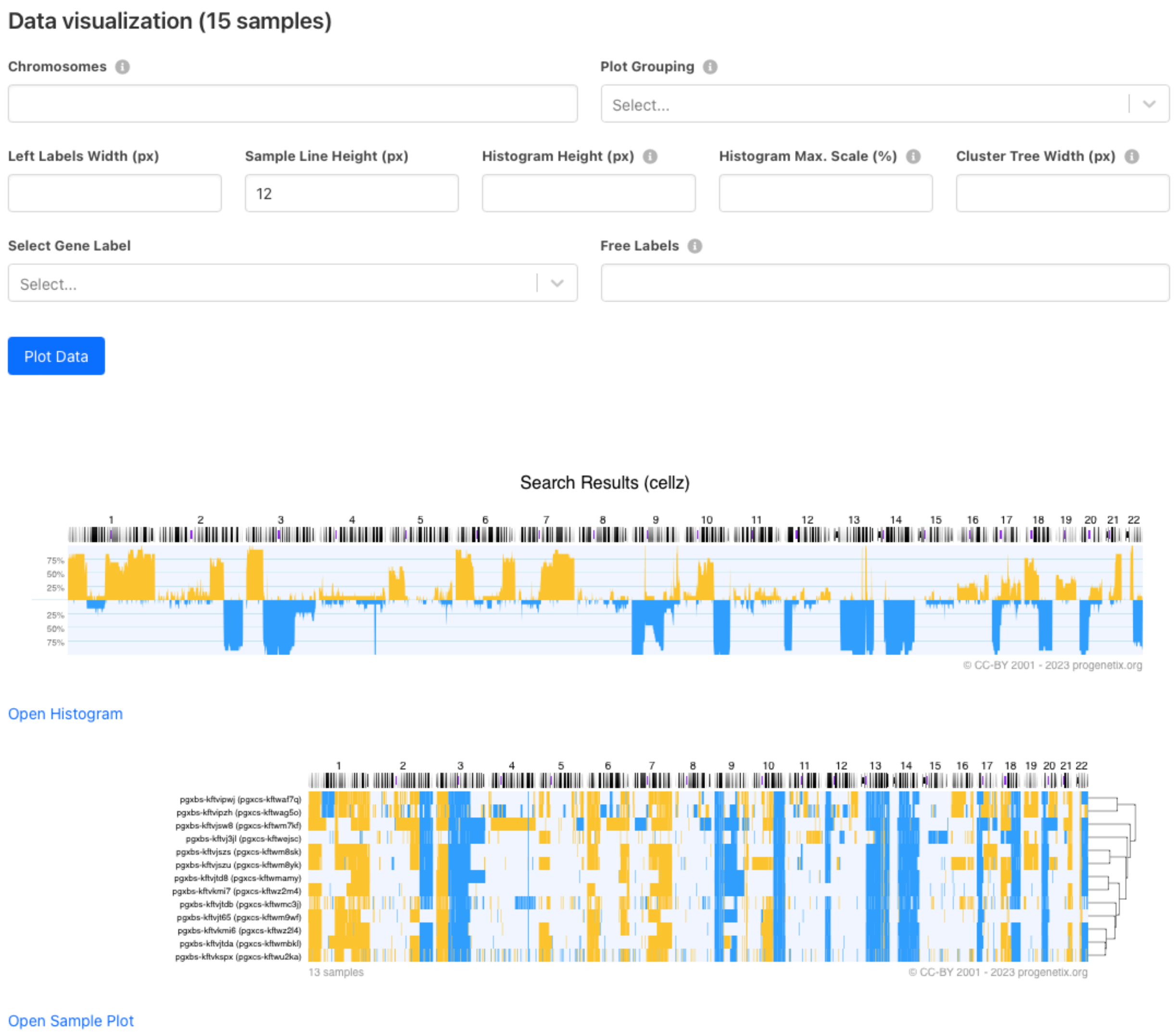


Visualization options land on a separate page with advanced options for data visualization. In addition to frequency plot, a sample plot with all clustered individual plots is shown.

Biosamples tab lists samples for this query. Associated diagnostic codes are also shown. Identifiers column shows other known sample identifiers such as GEO IDs or PMID.

Results Biosamples Biosamples Map

| Biosample Id | Dx Classifications | Identifiers |
|---|---|---|
| pgxbs-c43B6e8E | NCIT:C9110 Blast phase chronic myelogenous leukemia, BCR-ABL1 positive<br>pgx:icdom-98753 Chronic myelogenous leukemia, BCR/ABL positive<br>pgx:icdot-C42.1 Bone marrow<br>UBERON:0002371 bone marrow | cellosaurus:CVCL_0004 K-562 |
| pgxbs-6B0f6761 | NCIT:C9110 Blast phase chronic myelogenous leukemia, BCR-ABL1 positive<br>pgx:icdom-98753 Chronic myelogenous leukemia, BCR/ABL positive<br>pgx:icdot-C42.1 Bone marrow<br>UBERON:0002371 bone marrow | cellosaurus:CVCL_3827 K562/Adr |
| pgxbs-kftvipwj | NCIT:C9110 Blast phase chronic myelogenous leukemia, BCR-ABL1 positive<br>pgx:icdom-98753 Chronic myelogenous leukemia, BCR/ABL positive<br>pgx:icdot-C42.1 Bone marrow<br>UBERON:0002371 bone marrow | PMID:16001072 Garraway LA, Widlund HR et al. (2005): Integrative genomic analyses identify MITF as...<br>cellosaurus:CVCL_0004 K-562<br>geo:GSM50195<br>geo:GSE2520<br>geo:GPL2014 |
| pgxbs-kftvipzh | NCIT:C9110 Blast phase chronic myelogenous leukemia, BCR-ABL1 positive<br>pgx:icdom-98753 Chronic myelogenous leukemia, BCR/ABL positive<br>pgx:icdot-C42.1 Bone marrow<br>UBERON:0002371 bone marrow | PMID:16001072 Garraway LA, Widlund HR et al. (2005): Integrative genomic analyses identify MITF as...<br>cellosaurus:CVCL_0004 K-562<br>geo:GSM50259<br>geo:GSE2520<br>geo:GPL2015 |
| pgxbs-kftvj3jl | NCIT:C9110 Blast phase chronic myelogenous leukemia, BCR-ABL1 positive<br>pgx:icdom-98753 Chronic myelogenous leukemia, BCR/ABL positive<br>pgx:icdot-C42.1 Bone marrow<br>UBERON:0002371 bone marrow | PMID:20215515 Rothenberg SM, Mohapatra G et al. (2010): A genome-wide screen for microdeletions reveals...<br>cellosaurus:CVCL_3827 K562/Adr<br>geo:GSM827221<br>geo:GSE20306<br>geo:GPL3720 |

Results Biosamples Biosamples Map

Sample Origins

The map represents the origins of the matched samples, as derived from the original publication or resource repository. In the majority of cases this will correspond to the proxy information of the corresponding author's institution. Additional information can be found in the Geographic Coordinates documentation.

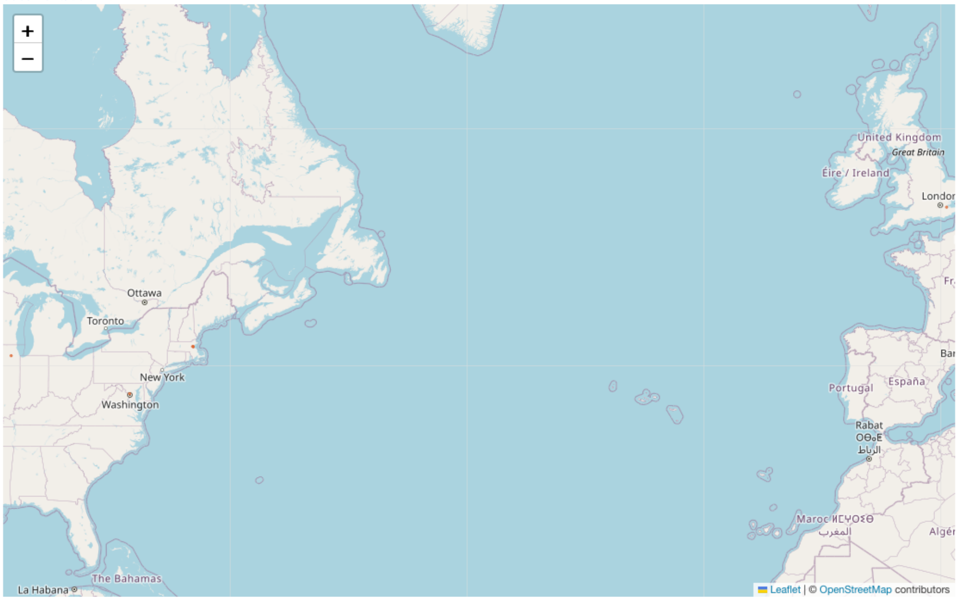

We have geographical info about where the sample was processed. That can be visualized in the Biosamples Map tab.

#### Search Results: SNV query

To access our SNV data, cell line on interest can be queried in "Search Cell Lines" like for CNV samples but additional field needs to be entered under Query by Position -> Variant Type -> SO:0001059 (any sequence alteration - SNV, INDEL…). This yields in all available SNVs for the cell line of interest. For example, search results for PC-9 cell line SNVs (Filtering Options -> only match exact terms) result in:

cellz

**Matched Samples:** 1
**Retrieved Samples:** 1

Dataset Responses (JSON)

Visualization options

Results | Biosamples | Variants

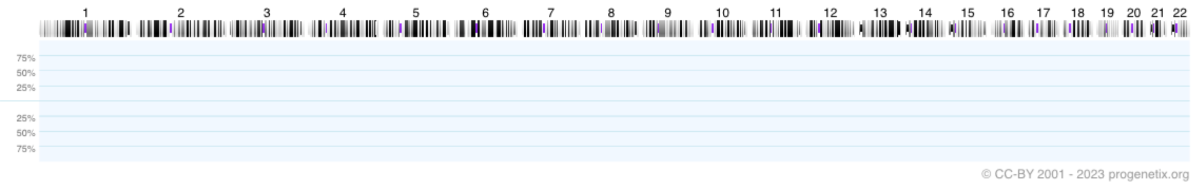


Reload histogram in new window

| Matched Subset Codes | Subset Samples | Matched Samples | Subset Match Frequencies |
|---|---|---|---|
| cellosaurus:CVCL_B260 | 68 | 1 | 0.015 |
| NCIT:C3512 | 1773 | 1 | 0.001 |

CNV plot is empty in this case as no CNV samples were included in the query. Variants are listed under "Variants" tab. Here, a small part of matched variants is listed. The results can be sorted by Digest, Gene, Pathogenicity or Variant Effect by clicking on the column name. Click on the variant ID in blue for more information on the variant.

Results | Biosamples | Variants

| Digest | Gene | Pathogenicity | Variant Effect | Variant Instances |
|---|---|---|---|---|
| 17:7674219-7674220:C>T | TP53 | Pathogenic | | V: pgxvar-63ce6a923319d2172d27fa18<br>B: pgxbs-9BCf68fA<br>V: pgxvar-63ce6adba24c83054b917026<br>B: pgxbs-9BCf68fA |
| 7:55174771-55174786:AGGAATTAAGAGAAGC>A | EGFR | Conflicting interpretations of pathogenicity | | V: pgxvar-644fd17a72c0f659d200c6c8<br>B: pgxbs-9BCf68fA |
| 5:35646791-35646792:G>A | SPEF2 | | Silent mutation | V: pgxvar-63ce6adba24c83054b917196<br>B: pgxbs-9BCf68fA |
| 2:21016551-21016552:G>T | APOB | | Silent mutation | V: pgxvar-63ce6adba24c83054b9170c0<br>B: pgxbs-9BCf68fA |

The example below shows available data for a variants collected from CCLE mutation set.

#### Variant Details for *pgxvar-63ce6adba24c83054b9170c0* (cellz)

##### Variant Shorthand Expression

- 2:21016551-21016552:G>T

##### Sample Information

- Sample: pgxbs-9BCf68fA

##### Molecular Attributes

- Gene: **APOB**
- Molecular effect: Silent mutation
- Aminoacid changes:
  - L1073L

##### Variant Identifiers

- Genomic HGVSID:

  g.chr2:21016552G>T

##### Variant Alternative IDs

- Ensembl:ENST00000233242.1 - Ensembl

##### Annotation Sources

- CCLE mutations

Here is an example for a matched variant in ClinVar dataset :

##### Variant Details for *pgxvar-63ce6a923319d2172d27fa18* (cellz)

**Variant Shorthand Expression**

- 17:7674219-7674220:C>T

**Sample Information**

- Sample: pgxbs-9BCf68fA

**Molecular Attributes**

- Gene: **TP53**
- Aminoacid changes:
  - R248Q
  - R116Q
  - R209Q
  - R89Q

**Variant Identifiers**

- Protein HGVSIDs:
  - P04637:p.Gly245Ser
- Genomic HGVSID:

  NC_000017.11:g.7674230C>T
- ClinVar IDs:
  - VCV000012356

**Variant Alternative IDs**

- ClinGen:CA000387 - ClinGen
- UniProtKB:P04637#VAR_005983 - UniProtKB
- OMIM:191170.0010 - OMIM
- dbSNP:11540652 - dbSNP

**Clinical Interpretations**

Clinical Relevance: **Pathogenic**

| ID | Description | Additional Annotations |
|---|---|---|
| MONDO:0006485 | Uterine carcinosarcoma | MedGen:C0280630, MONDO:0006485 |
| MONDO:0005082 | Prostate adenocarcinoma | MedGen:C0007112, MONDO:0005082 |
| MedGen:C0153574 | Malignant neoplasm of body of uterus | MedGen:C0153574 |
| MONDO:0006046 | Ovarian serous cystadenocarcinoma | MedGen:C0279663, MONDO:0006046 |

### CNV Profiles by Cancer Type

Like in Cell Line Listings, diagnostic codes are organized hierarchically. We provide NCIT and ICDO diagnostic codes for cancer cell lines. Number of biosamples and CNV profiles per NCI term are indicated. Clicking on available samples will lead to search form that can be used as described above.
